# DTI reveals whole-brain microstructural changes in the P301L mouse model of tauopathy

**DOI:** 10.1101/2020.10.28.358465

**Authors:** Aidana Massalimova, Ruiqing Ni, Roger M. Nitsch, Marco Reisert, Dominik von Elverfeldt, Jan Klohs

## Abstract

**Introduction:** Increased expression of hyperphosphorylated tau and the formation of neurofibrillary tangles are associated with neuronal loss and white matter damage. Using high resolution *ex vivo* diffusion tensor imaging (DTI), we investigated microstructural changes in the white and grey matter in the P301L mouse model of human tauopathy at 8.5 months-of-age. For unbiased computational analysis, we implemented a pipeline for voxel-based analysis (VBA) and atlas-based analysis (ABA) of DTI mouse brain data.

**Methods:** Hemizygous and homozygous transgenic P301L mice and non-transgenic littermates were used. DTI data were acquired for generation of fractional anisotropy (FA), mean diffusivity (MD), radial diffusivity (RD), axial diffusivity (AD) maps. VBA on the entire brain were performed using SPM8 and SPM Mouse toolbox. Initially, all DTI maps were co-registered with Allen mouse brain atlas to bring them to one common coordinate space. In VBA, co-registered DTI maps were normalized and smoothed in order to perform two-sample t-tests to compare hemizygotes with non-transgenic littermates, homozygotes with non-transgenic littermates, hemizygotes with homozygotes on each DTI parameter map. In ABA, the average values for selected regions-of-interests were computed with co-registered DTI maps and labels in Allen mouse brain atlas. After, the same two-sample t-tests were executed on the estimated average values.

**Results:** We made reconstructed DTI data and VBA and ABA pipeline publicly available. With VBA, we found microstructural changes in the white matter in hemizygous P301L mice compared to non-transgenic littermates. In contrast, more pronounced and brain-wide spread changes were observed in VBA when comparing homozygous P301L mice with non-transgenic littermates. Statistical comparison of DTI metrics in selected brain regions by ABA corroborated findings from VBA. FA was found to be decreased in most brain regions, while MD, RD and AD were increased compared to hemizygotes and non-transgenic littermates.

**Discussion/Conclusion:** High resolution *ex vivo* DTI demonstrated brain-wide microstructural changes in the P301L mouse model of human tauopathy. The comparison between hemizygous and homozygous P301L mice revealed a gene-dose dependent effect on DTI metrics. The publicly available computational data analysis pipeline can provide a platform for future mechanistic and longitudinal studies.

## Introduction

Alzheimer’s disease (AD), progressive supranuclear palsy, and corticobasal degeneration and frontotemporal dementia including frontotemporal dementia with Parkinsonism linked to chromosome 17 (FTDP-17) and Pick’s disease are characterized by the progressive formation of neurofibrillary tangles (NFTs) [1, 2]. NFTs are intracellular neuronal lesions composed of insoluble, conformationally abnormal, hyperphosphorylated accumulations of tau in neurons and glia [2]. The presence of NFTs in the brain triggers a complex cascade of biochemical and cellular processes that result in neurodegeneration and neuronal loss [2]. The occurrence of NFTs correlates well with spatial patterns of neuronal loss, and is related to the degree of cognitive deficits [3].

Magnetic resonance imaging (MRI) has been used to non-invasively characterize tissue changes in the brain associated with tau pathology. Structural MRI studies have shown that NFT deposition is associated with grey matter atrophy that results from neurodegeneration and that different tauopathies display distinctive regional patterns of grey matter atrophy [4-7]. In addition, white matter changes such as white matter atrophy and the occurrence of white matter hyperintensities have been reported [8]. Such observations are indicative of white matter damage and demyelination caused by axonal tau pathology [9]. Despite, their usefulness structural MRI provides little information about the underlying neurodegenerative changes in the brain.

Diffusion-weighted imaging is a MRI technique that probes the directional diffusivity of water molecules and can thus yield information about the microstructural properties of brain tissue [10-12]. Diffusion imaging data are commonly analyzed using a tensor model, i.e. applying diffusion tensor imaging (DTI) [13]. DTI shows sensitivity and specificity to axon and myelin pathology, where breakdown of white matter integrity results in measurable differences in diffusion of water molecules [14, 15]. DTI has been widely used to assess white matter abnormalities related to tau pathologies [5, 16-22]. A few DTI studies have also reported changes in diffusivity in the grey matter [23, 24]. Importantly, DTI microstructural changes seem to precede gross anatomical changes seen on structural MRI [23].

Genetic studies show the development of neurodegenerative tauopathies is associated with missense mutations in the microtubule associated tau protein (MAPT) [25, 26]. Based on this finding, transgenic mouse lines with MAPT mutations gene including P301S [27] and P301L [28-31] have been developed. Some models have the advantage of tetracycline-controlled transcriptional suppression of tau [31]. Histological studies have demonstrated deposition of neurofibrillary tangles pathology in particular in the cortex and hippocampus and marked atrophy of these areas [27-32]. Behavioral analysis revealed cognitive deficits in learning and motor tasks [30, 31].

Transgenic mouse models of tauopathy have been used in MRI studies [33-38]. Previous DTI studies assessing microstructural changes in rTg4510 mice, which express transgenic tau P301L mutations driven by a Ca^2+^/calmodulin kinase II (CaMKII) promoter system [30, 31] in the white and grey matter produced varied results [33-36]. Thus, we wished to determine DTI metrics in transgenic P301L mice [28, 29]. In the present study, we used *ex vivo* DTI to assess microstructural changes in 8.5-month old P301L mice in both white and grey matter regions. We leveraged the ability of long data acquisitions that are possible in *ex vivo* DTI to achieve a high spatial resolution, without suffering from image corruption due to motion or breathing that are common *in vivo* DTI [11, 12]. We implemented a pipeline for voxel-based analysis (VBA) and atlas-based analysis (ABA) for unbiased computational analysis of DTI mouse brain data. To test for gene-dose-dependent effects, we assessed both hemizygous and homozygous P301L mice and compared them with non-transgenic littermates. We hypothesized that effects on DTI microstructural parameters would be stronger in homozygotes than in hemizygotes, because of the stronger tau transgene expression in homozygous P301L mice.

## Materials and Methods

### Animal Models

Hemizygous mice (n=8; 5 females/3 males), homozygous (n=8; 5 females/3 males) transgenic for a human 4 repeat tau isoform with the P301L under Thy 1.2 promoter of 8.5 months of age and age-matched non-transgenic littermates 8.5 (n=8; 2 females/6 males) were used. Animals were housed in ventilated cages inside a temperature-controlled room, under a 12-hour dark/light cycle. Each cage housed up to five mice. Paper tissue and red Tecniplast mouse house^®^ (Tecniplast, Milan, Italy) shelters were placed in cages as environmental enrichments. Pelleted food (3437PXL15, CARGILL) and water were provided ad libitum.

### Magnetic Resonance Imaging

Mice were intracardially perfused under deep anesthesia (ketamine/xylazine/acepromazine maleate (75/10/2 mg/kg body weight, i.p. bolus injection) with 0.1 M PBS (pH 7.4). Heads were removed post-fixed in 4 % paraformaldehyde in 0.1 M PBS (pH 7.4) for 6 days and stored in 0.1 M PBS (pH 7.4) at 4°C. Brains were not removed from the skull, which has been shown previously to preserve cortical and central brain structure [35]. The heads were placed in a 15 ml centrifuge tube filled with perfluoropolyether (Fomblin Y, LVAC 16/6, average molecular weight 2700, Sigma-Aldrich, U.S.A.). Data was acquired on a small animal MRI system (BioSpec 94/30 animal MRI system, Bruker Corporation) at 400 MHz and equipped with a BGA 12AS HP gradient system with a maximum gradient strength of 400 mT/m and minimum rise time of 70 μs. A cryogenic 2 x 2 radio frequency surface coil probe (overall coil size 20×27 mm^2^, Bruker BioSpin AG, Fällanden, Switzerland) with the coil system operating at 30 K (Bruker BioSpin AG, Fällanden, Switzerland) was used in combination with a circularly polarized 86 mm volume resonator for transmission.

DTI was acquired with 3D multi-shot echo planar imaging (EPI) sequence (4 shots) with a field of view of 18 mm x 12 mm x 9 mm, matrix dimension of 180 x 120 x 90, resulting in a nominal voxel resolution of 100 μm x 100 μm x 100 μm. The following imaging parameters were chosen: a repetition time of 1500 ms, an echo time of 28 ms, no averaging, 5 volumes acquired with a b values of 0 (A0), b-value of 4000 s/mm^2^ along 60 diffusion encoding directions. A global and MAPSHIM protocol with a field map (default settings) was used for shimming. The total acquisition time per brain was 9 h 45 minutes.

### Image Processing

Image processing was performed with the Nora medical imaging platform (http://www.nora-imaging.com/; University Medical Center Freiburg, Germany) [39, 40]. All the data was first uploaded to the platform and automatically converted into NIFTi format.

Diffusion tensors were estimated using DTI&Fibertools (https://www.uniklinik-freiburg.de/mr-en/research-groups/diffperf/fibertools.html) and on this basis fractional anisotropy (FA), mean diffusivity (MD), radial diffusivity (RD), axial diffusivity (AD) maps were computed. To normalize images to atlas space SPM8 (Wellcome Trust Centre for Neuroimaging, UCL, UK) together with the SPM Mouse toolbox (University of Oxford, UK) was used, For each scan the B0 image was normalized to SPM’s mouse atlas space using SPM’s normalize and segment tool. Further the Allen mouse brain atlas (http://mouse.brain-map.org/static/atlas) was also nonlinearly registered to SPM’s atlas space to enable the voxel and atlas based analysis. All considered microstructural maps (FA, MD, RD, AD) were finally warped to atlas space for VBA and ABA.

### Voxel-Based Analysis

VBA on the entire brain were performed using SPM8. In addition, the SPM Mouse toolbox (University of Oxford, UK) was used, because it extends the SPM functionality to the animal brains. All normalized images were smoothed (0.1 x 0.1 x 0.1 mm^2^ Gaussian) before performing a two-sample t-test with a significance level of p<0.05. Besides, the average maps of each parameter for every subgroup were generated to visualize the estimated t-values on it. In total 3 two-sample t-tests were executed including the comparisons of hemizygotes vs. homozygotes, hemizygotes vs. non-transgenic littermates, homozygotes vs. non-transgenic littermates.

### Atlas-Based Analysis

Allen mouse brain atlas was overlaid on each normalized image. The atlas contains 670 labeled regions. Then, the average values of each region-of-interests (ROIs) from the atlas were computed. In order to limit the number of candidate regions for the examination, we selected anterior commissure (AC), corpus callosum (CC), cerebral peduncle (CPD), cortex (CRX), hippocampal region (HPR), subiculum (SUB) and thalamus (TH). An one-way analysis of variance (ANOVA) was used to determine differences between AD, MD, RD, FA in these ROIs between hemizygotes vs. homozygotes, hemizygotes vs. non-transgenic littermates, homozygotes vs. non-transgenic littermates. We used p<0.05, p<0.01 and p<0.0001 as our significance level. All computations were done using Python version 3.5.0.

## Results

The reconstructed data is available in the repository (DOI: xxx). We also made the code for VBA () and ABA available (https://github.com/aidanamv/ABA-pipeline).

We examined *ex vivo* DTI metrics of the brains of hemizygous and homozygous P301L mice at 8.5 months of age. Resulting maps of the FA, MD, RD and AD are shown in Figure 1 and in different views in Supplementary Figure 1-4. The high-resolution FA maps show clearly white matter structures that are highly anisotropic, like the corpus callosum and the anterior and hippocampal commissure.

**Fig. 1.**
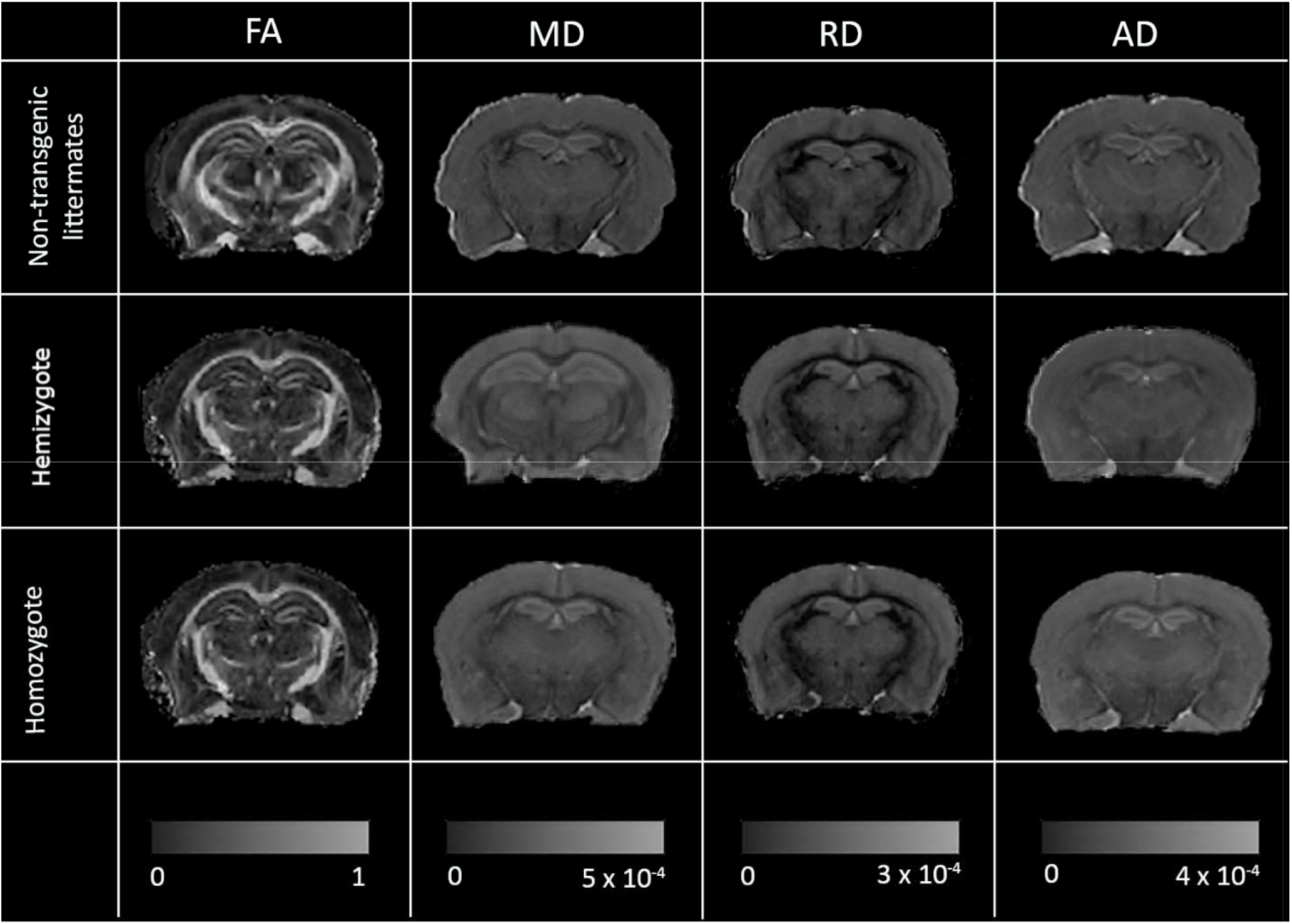
Representative coronal FA, MD, RD and AD maps of 8.5-months old non-transgenic littermates, hemizygous and homozygous P301L mice. Shown are views at approximately −1.8 mm from Bregma. AD, RD, and MD values are in units of ×10^−4^ mm^2^/s.

### VBA reveals gene-dose dependent effects on DTI metrics

VBA was performed to compare differences in FA, MD, RD and AD between groups and to test for gene-dose dependent effects. First, we compared hemizygous P301L mice with non-transgenic littermates (shown in Fig. 2). The color gradient represents the magnitude of significant differences between hemizygous mice compared to the non-transgenic littermate group (two-sample t-test). We observed minor changes in FA and AD in white matter structures such as the corpus callosum and hippocampal commissure. In contrast, more pronounced changes in the white matter in all DTI metrics were observed in the VBA when comparing homozygote P301L mice with non-transgenic littermates (shown in Fig. 3). In addition, to the changes seen in white matter structures, group differences were also detected in grey matter structures such as the cortex, hippocampus, and thalamus for all DTI metrics. Subsequently, we compared hemizygotes with homozygotes (shown in Fig. 4). Group differences were seen in all DTI metrics in the hippocampal commissure only.

**Fig. 2.**
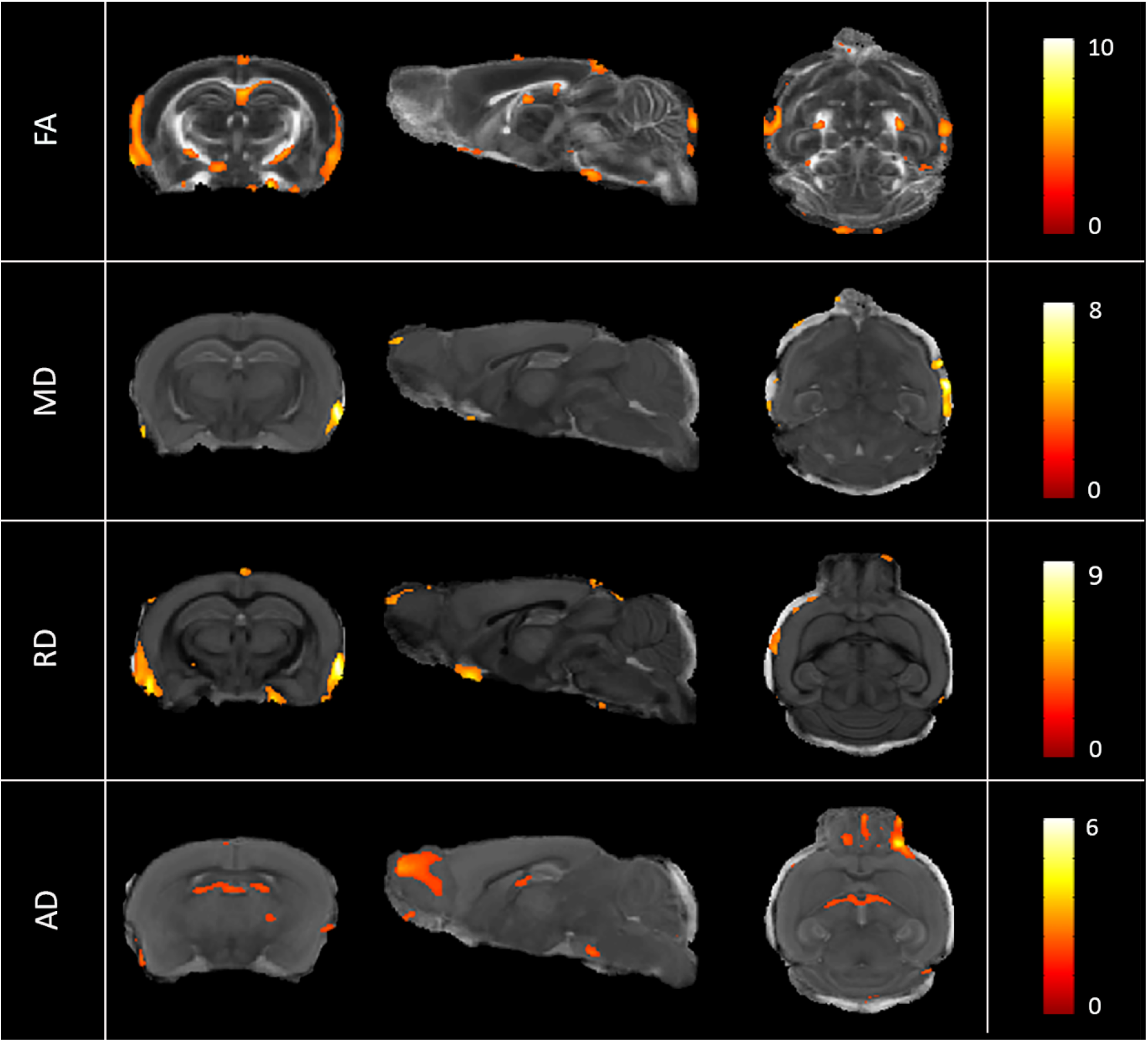
Mild DTI alterations in hemiyzygous P301L mice. VBA (color-coded t-values overlaid on an anatomical mouse brain template) comparing hemiyzygous P301L mice with non-transgenic littermates (p<0.05, uncorrected). Changes in DTI metrics were mainly observed in the white matter (e.g. hippocampal commissure).

**Fig. 3.**
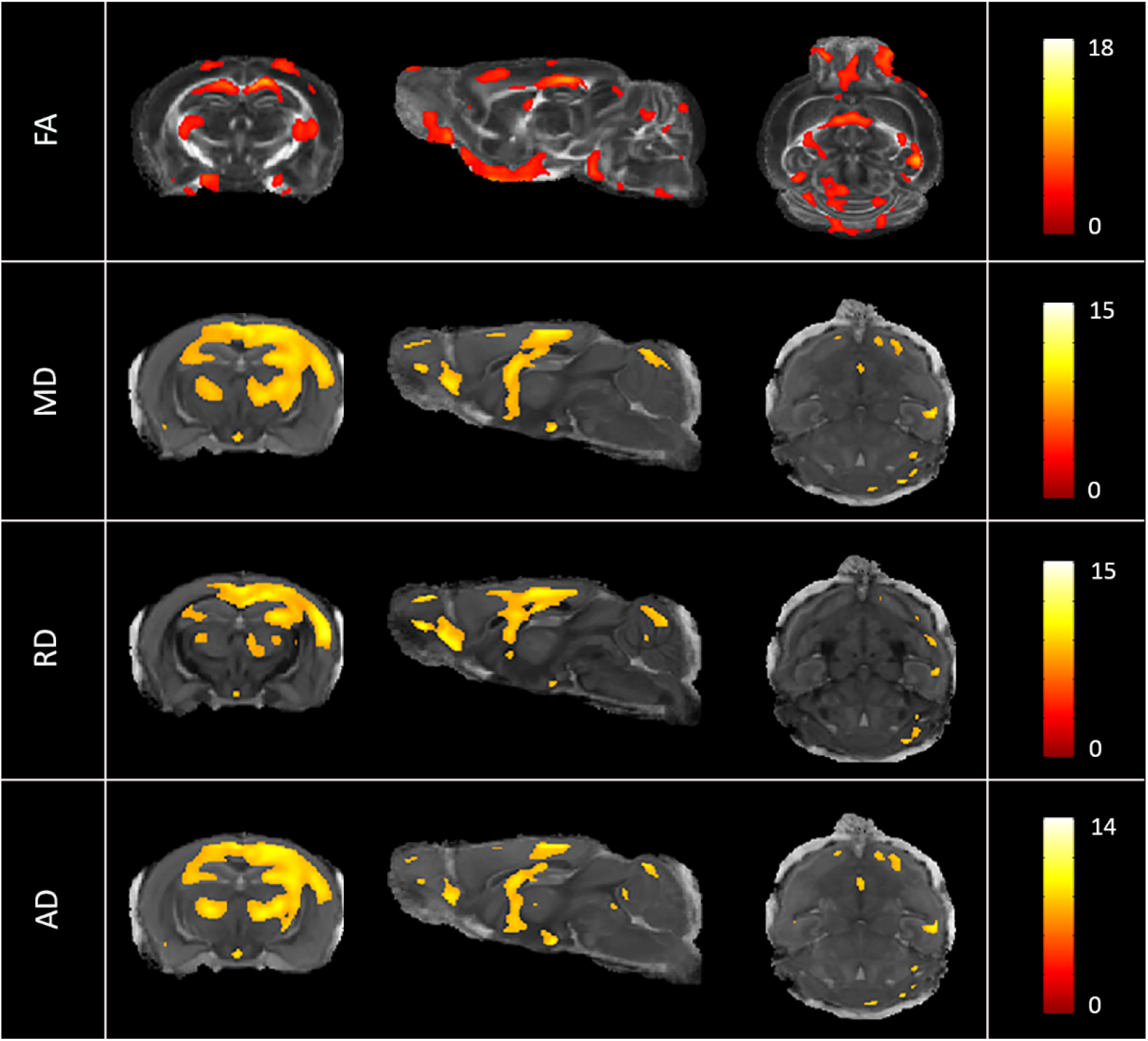
Severe DTI alterations in homozygous P301L mice. VBA (t-values overlaid on mouse brain template) comparing homozygous P301L mice with non-transgenic littermates (p<0.05, uncorrected). Changes in DTI metrics were observed across the brain, in both grey and white matter areas.

**Fig. 4.**
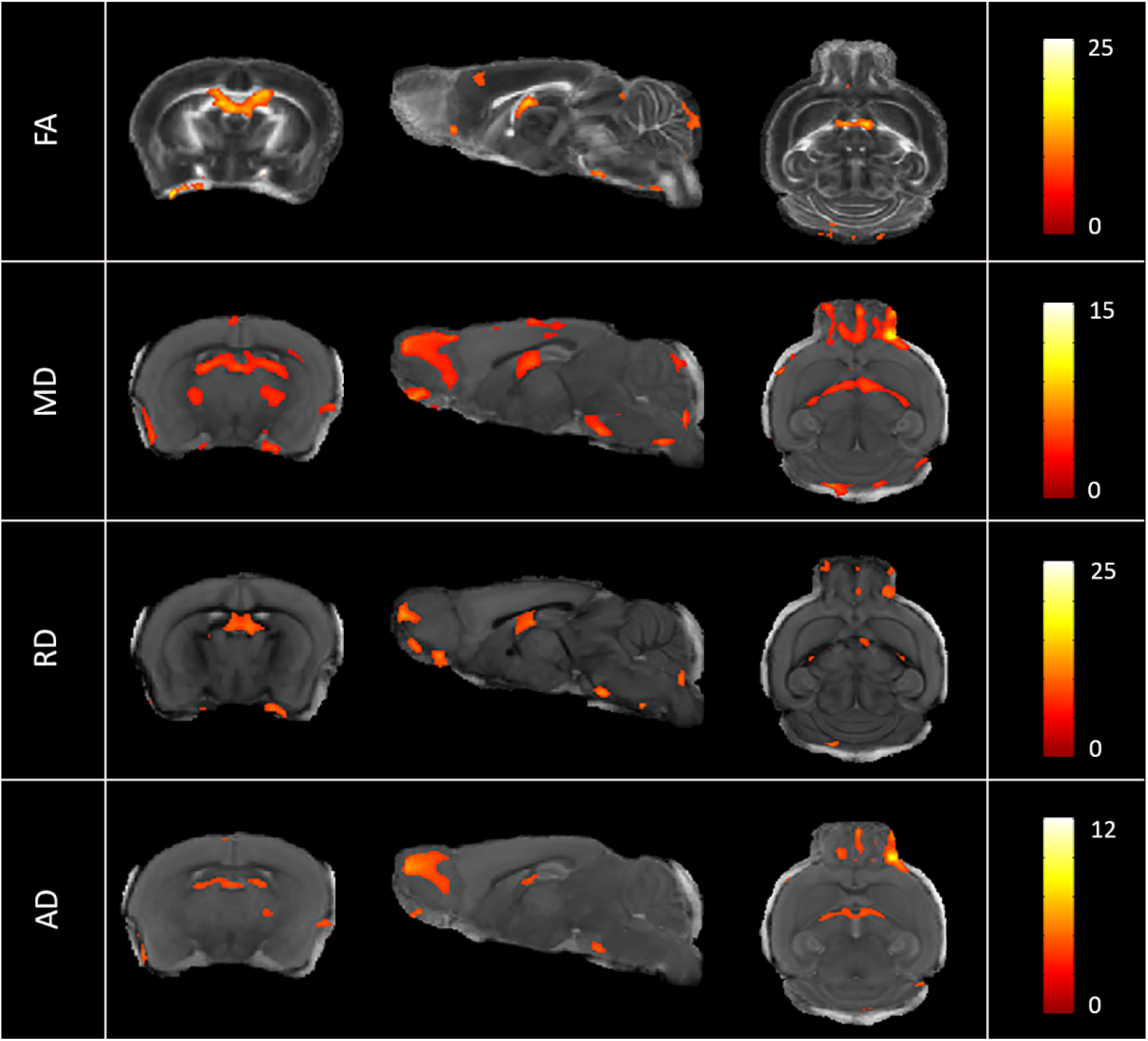
Grade of tau gene expression determine severity of DT alterations. VBA (t-values overlaid on mouse brain template) comparing hemizygous with homozygous P301L mice (p<0.05, uncorrected). Changes in DTI metrics were observed across the brain.

### ABA reveals grey and white matter alterations related to tau pathology

The results of the ABA analysis are shown in Table 1 and Figure 5. We analyzed selected white matter structures and grey matter areas (Fig. 5). Statistical analysis was performed in three white matter regions (anterior commissure, corpus callosum and cerebral peduncle) and four grey matter regions (hippocampal regions, cortex, subiculum and thalamus).

**Table 1.** Diffusion tensor imaging parameters for different regions from atlas based analysis.

| Region | DTI parameter | Non-transgenic littermates (n=8) | Hemizygous P301L (n=8) | Homozygous P301L (n=8) |
| --- | --- | --- | --- | --- |
| Anterior commissure | FA | 0.56±0.08 | 0.56±0.09 | 0.50±0.04 |
|  | MD | 0.14±0.01 | 0.15±0.03 | 0.19±0.02<sup>a,b</sup> |
|  | RD | 0.07±0.02 | 0.08±0.02 | 0.11±0.01<sup>a,b</sup> |
|  | AD | 0.18±0.01 | 0.19±0.04 | 0.23±0.01<sup>a,b</sup> |
| Corpus callosum | FA | 0.58±0.05 | 0.58±0.06 | 0.51±0.02<sup>a,b</sup> |
|  | MD | 0.13±0.01 | 0.15±0.03 | 0.18±0.01<sup>a,b</sup> |
|  | RD | 0.07±0.01 | 0.07±0.02 | 0.10±0.01<sup>a,b</sup> |
|  | AD | 0.17±0.02 | 0.15±0.03 | 0.17±0.01<sup>a,b</sup> |
| Cerebral peduncle | FA | 0.55±0.05 | 0.61±0.08 | 0.48±0.07<sup>a,b</sup> |
|  | MD | 0.18±0.02 | 0.16±0.03 | 0.21±0.02<sup>a,b</sup> |
|  | RD | 0.10±0.03 | 0.06±0.03 | 0.12±0.02<sup>b</sup> |
|  | AD | 0.23±0.02 | 0.21±0.03 | 0.25±0.02<sup>a,b</sup> |
| Cortex | FA | 0.26±0.02 | 0.28±0.02 | 0.24±0.02<sup>b</sup> |
|  | MD | 0.19±0.01 | 0.21±0.05 | 0.26±0.02<sup>a,b</sup> |
|  | RD | 0.15±0.01 | 0.15±0.04 | 0.20±0.01<sup>a,b</sup> |
|  | AD | 0.22±0.01 | 0.24±0.06 | 0.28±0.02<sup>a,b</sup> |
| Hippocampal region | FA | 0.28±0.02 | 0.27±0.03 | 0.24±0.03<sup>a,b</sup> |
|  | MD | 0.22±0.01 | 0.23±0.04 | 0.28±0.02<sup>a,b</sup> |
|  | RD | 0.17±0.01 | 0.18±0.04 | 0.22±0.02<sup>a,b</sup> |
|  | AD | 0.25±0.01 | 0.26±0.05 | 0.30±0.02<sup>a,b</sup> |
| Subiculum | FA | 0.26±0.03 | 0.23±0.01 | 0.21±0.02<sup>a,b</sup> |
|  | MD | 0.19±0.01 | 0.20±0.05 | 0.23±0.03<sup>a</sup> |
|  | RD | 0.14±0.01 | 0.15±0.04 | 0.19±0.02<sup>a,b</sup> |
|  | AD | 0.21±0.01 | 0.23±0.06 | 0.26±0.03<sup>a</sup> |
| Thalamus | FA | 0.33±0.04 | 0.32±0.04 | 0.27±0.03<sup>a,b</sup> |
|  | MD | 0.16±0.01 | 0.17±0.04 | 0.21±0.02<sup>a,b</sup> |
|  | RD | 0.11±0.01 | 0.12±0.03 | 0.16±0.02<sup>a,b</sup> |
|  | AD | 0.19±0.01 | 0.20±0.05 | 0.23±0.02<sup>a</sup> |

mean±SD; AD, RD, and MD values are in units of $\times 10^{-4}$ mm<sup>2</sup>/s; <sup>a</sup> p< 0.05 vs non-transgenic littermates; <sup>b</sup> p< 0.05 vs hemizygous. One-way ANOVA.

**Fig. 5.**
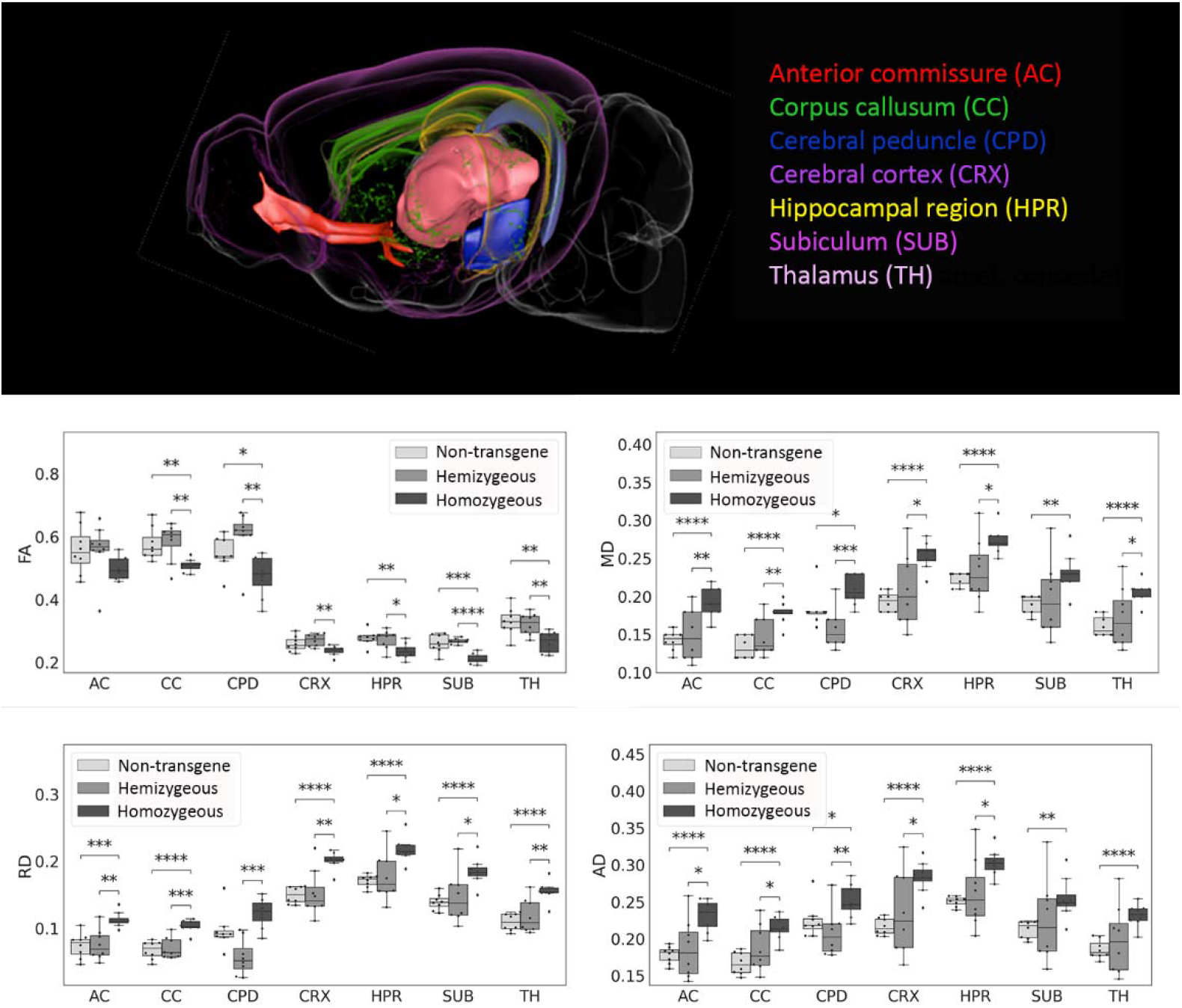
Gene-dose dependent effects on DTI metrics. ABA of selected brain regions in non-transgenic littermates, hemizygous and homozygous P301L mice. Regions analysed where the anterior commissure (AC), corpus callosum (CC), cerebral peduncle (CPD), cerebral cortex (CRX), hippocampal region (HPR), subiculum (SUB), and thalamus (TH). Data are shown as mean ± standard deviation. Oneway ANOVA. *p<0.05, ** p<0.01, *** p<0.001.

### (i) Fractional anisotropy

In the anterior commissure, there was no difference in FA between groups observed. In the corpus callosum, cerebral peduncle, hippocampal regions, subiculum and thalamus FA values were significantly decreased in homozygous P301L mice compared to hemizygous transgenic and non-transgenic littermates. In the cortex, FA values were found reduced in homozygous compared to hemizygous transgenic P301L mice. There were no statistical differences between hemizygote P301L mice and non-transgenic controls.

### (ii) Mean diffusivity

MD values were significantly increased homozygote P301L mice compared to non-transgenic littermates in all regions investigated, except for the subiculum. Values were also significantly higher than in hemizygotes. There was no statistical differences in MD values between hemizygous P301L mice and non-transgenic controls.

### (iii) Radial diffusivity

RD values were found significantly increased in the anterior commissure, corpus callosum, cortex, hippocampal region, subiculum and thalamus in homozygous P310L mice compared to hemizygous transgenic mice and non-transgenic littermates. In the cerebral peduncle, RD values were found significantly increased in homozygous P301L mice compared to hemizygotes, but were not statistically different from non-transgenic littermates. No differences were observed between hemizygous P301L mice and non-transgenic littermates.

### (iv) Axial diffusivity

AD values were significantly found increased in all regions analysed in homozygous P301L mice compared to non-transgenic littermates. In the anterior commissure, corpus callosum, cerebral peduncle, cortex and hippocampal regions AD values were also significantly higher in homozygous P301L mice compared to hemizygous mice, while no differences were observed in the subiculum and thalamus. No differences were observed between hemizygous P301L mice and non-transgenic littermates in all regions tested.

## Discussion/Conclusion

In this study, we acquired *ex vivo* DTI data in 8.5-month old hemizygous and homozygous P301L mice and age-matched non-transgenic littermates. A pipeline for post-processing of mouse brain DTI data, and VBA and ABA for statistical analysis revealed microstructural changes in both white and grey matter regions. In homozygotes, FA was found decreased in most brain regions, while MD, RD and AD were increased compared to hemizygotes and non-transgenic littermates. We found changes in DTI metrics to be more pronounced in homozygotes than in hemizygotes.

DTI has been previously used in patient studies to assess microstructural changes related to tau pathologies [5, 16-24]. Application of DTI to animal models of tauopathy may be useful for validation and mechanistic studies and monitoring of tau-targeted therapies. Previously, DTI has been applied to characterize brain tissue changes in the rTg4510 strain [33-36], but produced varied results. Thus, we aimed to investigate the effects of tau pathology on DTI metrics in the P301L models of human tauopathy.

In contrast to previous studies where DTI in mice was performed *in vivo* [33-36], we performed *ex vivo* DTI. Compared to the *in vivo* setting, *ex vivo* DTI has the advantage that it is not hampered by motion artifacts e.g. breathing. It also allows for prolonged scanning times. We acquired 3D data with i.e. 100 μm isotropic resolution in 9 h 45 min. We measured complete head samples, which preserve the structural integrity of the brain [35] in a high field small-bore magnet with cryogenic radiofrequency coils [43] to obtain diffusion data with sufficient signal-to-noise ratio for automated VBA and ABA analysis.

*Ex vivo* DTI requires chemical fixation with formaldehyde to preserve the tissue. Nevertheless, the procedure alters the macroscopic and microscopic properties of tissue. For example, a study by Ma et al. [44] has shown that most of the white structures demonstrated significantly larger *ex vivo* volumes after fixation than *in vivo* volumes (i.e. internal capsule, and fimbria) except for the smallest white matter structure (e.g. anterior commissure), which was significantly smaller *ex vivo*. Conversely, grey matter structures (e.g. neocortex, thalamus, and hippocampus) decreased in volume. Fixation does alter the magnitude of water diffusion in the tissue, but measures of the directionality of diffusion are less affected [45, 46]. As samples are equally treated, the volume changes induced by fixation do not hamper advanced, unbiased computational approaches and allow detection of anatomical phenotypes in transgenic animals. The reduced diffusivity of fixed tissue requires strong gradients for strong diffusion weighting. These are commonly available in small-bore animal scanners. We used a b value of 4000 s/mm^2^, as recommended for *ex vivo* DTI [47].

Previous studies have used ROI analysis [33-36], while we opted for VBA and ABA for statistical analysis of DTI data. For ROI analysis selected areas are manually drawn, which is more time-consuming, requires a priori anatomical localization, and is prone to bias [48]. Moreover, in many neurodegenerative diseases, microstructural changes occur across the whole brain, and conventional ROI analysis may miss to detect changes in areas that are not selected [49]. VBA has been widely used in structural MRI as an unbiased approach to assess morphological changes of tissue [44, 50]. Voxel-by-voxel statistical comparison is performed once all brain images are normalized to a template space, with the assumption that each individual voxel represents the same anatomical location of the brain [41, 49, 51].

With VBA, we found marked changes when comparing homozygous P301L mice with non-transgenic littermates, while we observed only minor differences between hemizygous transgenic mice and non-transgenic littermates. The comparisons demonstrated that changes in DTI metrics are gene-dose dependent, where mice with the stronger tau transgene expression (i.e. homozygous P301L mice) show the more pronounced phenotype. This finding is in line with the results of the ABA where differences in DTI metrics were greater between homozygous P301L mice and non-transgenic littermates than between hemizygous P301L mice and non-transgenic littermates. In addition, VBA revealed only small differences between hemizygous and homozygous P301L mice and between hemizygous transgenic mice and non-transgenic littermates, which is corroborated by ABA.

The spatial normalization of the brains in VBA was based on scalar image based registration, incorporating 12-parameter affine transformation and diffeomorphic mapping, which is essentially done to match internal brain structures to the template. One may apply a low-pass filter to smooth normalized images; however, the fact, if the particular location is the same voxel in other images, is still questionable due to the potential atrophy in the brain [41]. Smoothing in DTI is also challenging due to thin tracts of white matter and heterogeneity of the FA maps. Nevertheless, this method is still in use in biomedical imaging due to its simplicity [50]. Besides, its statistical power is poor due to the high level of noise.

In comparison, ABA analysis is based on automatic segmentation of brain regions defined in the atlas after normalization [41, 51-53]. This provides high sensitivity to small and widely distributed changes, but it fails when the regions are not within the anatomical boundaries of the predefined atlas [53]. Results from ABA shown in Table 1 revealed a decrease in FA in the corpus callosum and cerebral peduncle. We observed for all selected white matter structures increased MD, RD and AD compared to non-transgenic littermates. An earlier study by Sahara et al. reported age-dependent changes in DTI metrics in rTg4510 mice [33]. A decrease in FA values was noted in the corpus callosum, anterior commissure, internal capsule, splenium, and fimbria of 8-months-old rTG4510 mice. RD was found increased in the corpus callosum, splenium and fimbria while MD and AD were unchanged. In a study by Colgan et al. found lower FA values and higher MD in the corpus callosum of 8.5-months-odl rTG4510 mice compared to non-transgenic littermates [36]. A study by Wells et al. reported increased RD values in the corpus callosum in rTG4510, while FA, MD and AD were not different from non-transgenic littermates [34].

In grey matter structures, we found decreased FA values e.g. the hippocampal regions, subiculum and thalamus in P301L mice. A previous study by Wells reported increased FA and MD in the cortex and hippocampus of rTG4510 mice [34]. In the thalamus, the MD was found increased, while the FA was unchanged. A study by Holmes et al. reported increased FA and MD in the cortex and hippocampus of rTG4510 mice [35]. In the thalamus FA was increased, while MD was not different. In a study by Colgan et al. MD was increased in the hippocampus and cortex [36]. FA was higher in the hippocampus, but was not different in the cortex. No change in FA and MD have been observed in the thalamus.

DTI studies in rTg4510 mice of similar ages yielded variable results [33-36] and thus it is unclear if DTI changes in rTG4510 mice reflect the human situation. Notwithstanding, the current pattern of decreased FA and increased MD, RD and AD values in P301L mice are in good agreement with DTI changes observed in human tauopathies such as progressive supranuclear palsy [20], frontotemporal dementia [21], mild cognitive impairment [23] and Alzheimer’s disease [17, 24]. However, differences in DTI metrics in cross-sectional studies in transgenic mouse models of tauopathy based on MAPT mutations may arise due to the type of mutation, promoter, transgene expression levels, genetic background of strain and housing conditions [54]. Furthermore, variability of results may be due to different analysis pipelines used. Given the variability of reported DTI results in tau mouse models measures should be taken to improve the reproducibility of data analysis outcomes. We suggest that future studies share reconstructed FA, MD, RD and AD maps and analysis code, which would enable other groups to perform an independent analysis with the data, to validate the code used and to enable of image-based meta-analysis. Finding converging patterns of microstructural changes or identifying different phenotypes between different tau mouse models would increase the translational value of preclinical DTI studies.

DTI is widely used to assess microstructural properties of white matter and DTI metrics FA, MD AD and RD are sensitive to different aspects of white matter pathology [55]. A decrease in FA in homozygous P301L mice indicates white matter damage with loss of directionality of water diffusion [55]. An increase in AD may indicate acute axonal damage as accumulation of neurofilaments may hinder diffusion parallel to axons [16, 17]. RD is sensitive to myelin injury, and an increase in RD may indicate increases of extracellular space [22, 33]. While white matter damage such as swollen axons, fragmentation, vacuolization and inflammation have been reported for the rTG4510 mouse model [33, 37], white matter changes in the P301L strain [28, 29] used in the current study have not been characterized.

DTI is increasingly used to assess microstructural changes in the grey matter [56]. For GM cellular underpinnings of altered diffusivity measures are not yet well understood. It has been speculated that a reduced FA most likely results from a change in tissue cytoarchitecture due to subtle small vessel alterations, and possibly gliosis [24]. Increased MD most likely results from loss of neurons and dendrites, which result in an increase in extracellular space and elevated water diffusivity within these regions [24]. Neuronal loss and gliosis have been both described to occur in some grey matter regions of P301L mice [29, 38]. Pronounced changes in DTI metrics have been observed in regions of strong tau expression such as the hippocampus and cortex [29, 38, 57]. However, in previous studies a correlation between DTI and tau burden in rTg4510 mice was not established [34, 36].

Future studies may seek to examine microstructural changes *in vivo* in a longitudinally designed study. For example, it would be important to determine the onset of changes in DTI and how this is related to gross anatomical changes and cognitive function. Studies in patients with mild cognitive impairment have shown that changes in DTI metrics in the hippocampus, a region involved in working memory formation, can predict cognitive decline and constitute a more sensitive predictor than changes in hippocampal volume [5, 23]. Simultaneous multi-slice MRI combined with parallel imaging would significantly reduce scan time for in vivo DTI, but is challenging in the mouse due to limited space and small coil size [12]. Moreover, the varying contributions of tau deposition and down-stream cellular processes to changes in water diffusivity in the white and grey matter should be elucidated.

In summary, we have shown brain-wide microstructural changes in the P301L mouse model of human tauopathy with DTI. The publicly available DTI data and computational VBA and ABA data analysis pipeline may serve as a platform for future longitudinal studies and to mechanistic studies that help to gain insights in the interpretation of DTI data in tauopathies as well as facilitate findings between different laboratories.

## Statements

## Acknowledgement

The authors acknowledge Daniel Schuppli at the Institute for Regenerative Medicine, University of Zurich for technical assistance.

## Statement of Ethics

All experiments were performed in accordance with the Swiss Federal Act on Animal Protection and were approved by the Cantonal Veterinary Office Zurich (permit number: ZH082/18). All procedures fulfilled the ARRIVE guidelines on reporting animal experiments.

## Author Contributions

The study was conceived and designed by Ruiging Ni and Jan Klohs. Roger M. Nitsch provided the P301L mouse model. Marco Reisert and Dominik von Elverfeldt provided the software platform for DTI reconstruction. Material preparation, data collection and analysis were performed by Aidana Massalimovsa, Ruiqing Ni and Marco Reisert. The first draft of the manuscript was written by Aidana Massalimovsa and Jan Klohs. All authors commented on previous versions of the manuscript. All authors read and approved the final manuscript.

---

Suppl Fig. 1. Coronal, sagittal and horizontal views of the fractional anisotropy (FA) of non-transgenic littermates, hemizygous and homozygous littermates.

Suppl Fig. 2. Coronal, sagittal and horizontal views of the mean diffusivity (MD) of non-transgenic littermates, hemizygous and homozygous littermates.

Suppl Fig. 3. Coronal, sagittal and horizontal views of the radial diffusivity (RD) of non-transgenic littermates, hemizygous and homozygous littermates.

Suppl Fig. 4. Coronal, sagittal and horizontal views of the axial diffusivity (AD) of non-transgenic littermates, hemizygous and homozygous littermates.

