## Supplementary figures and images for "DTI reveals whole-brain microstructural changes in the P301L mouse model of tauopathy"

### Suppl figures

SFig 1

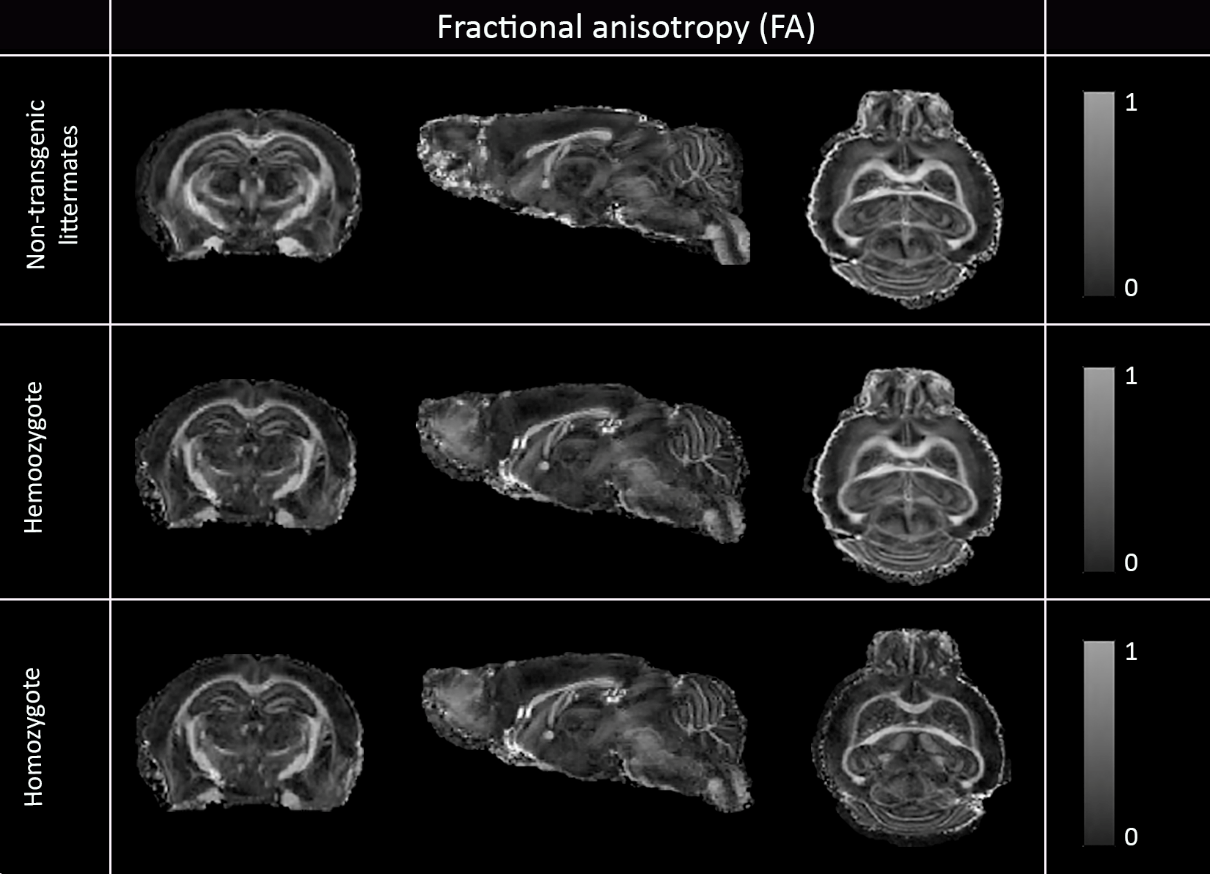

SFig 2

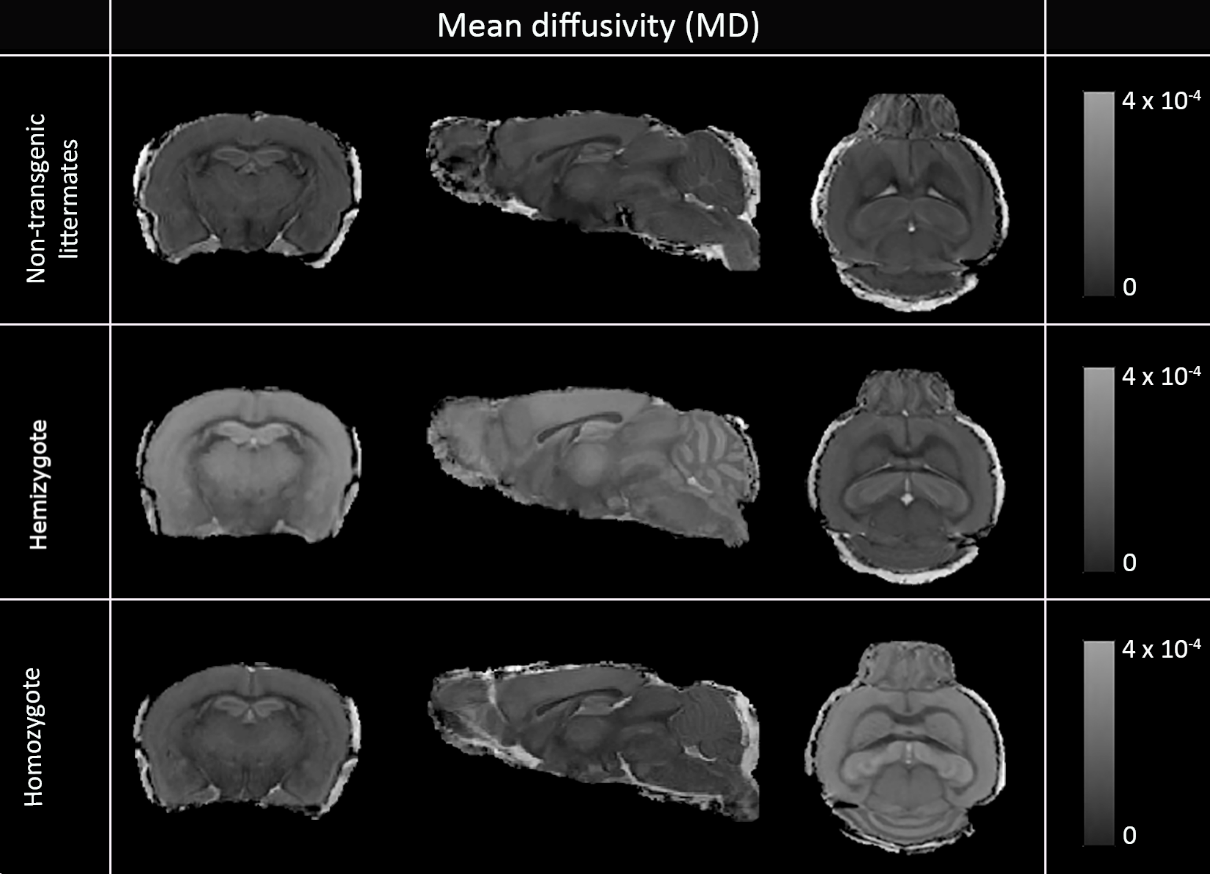

SFig 3

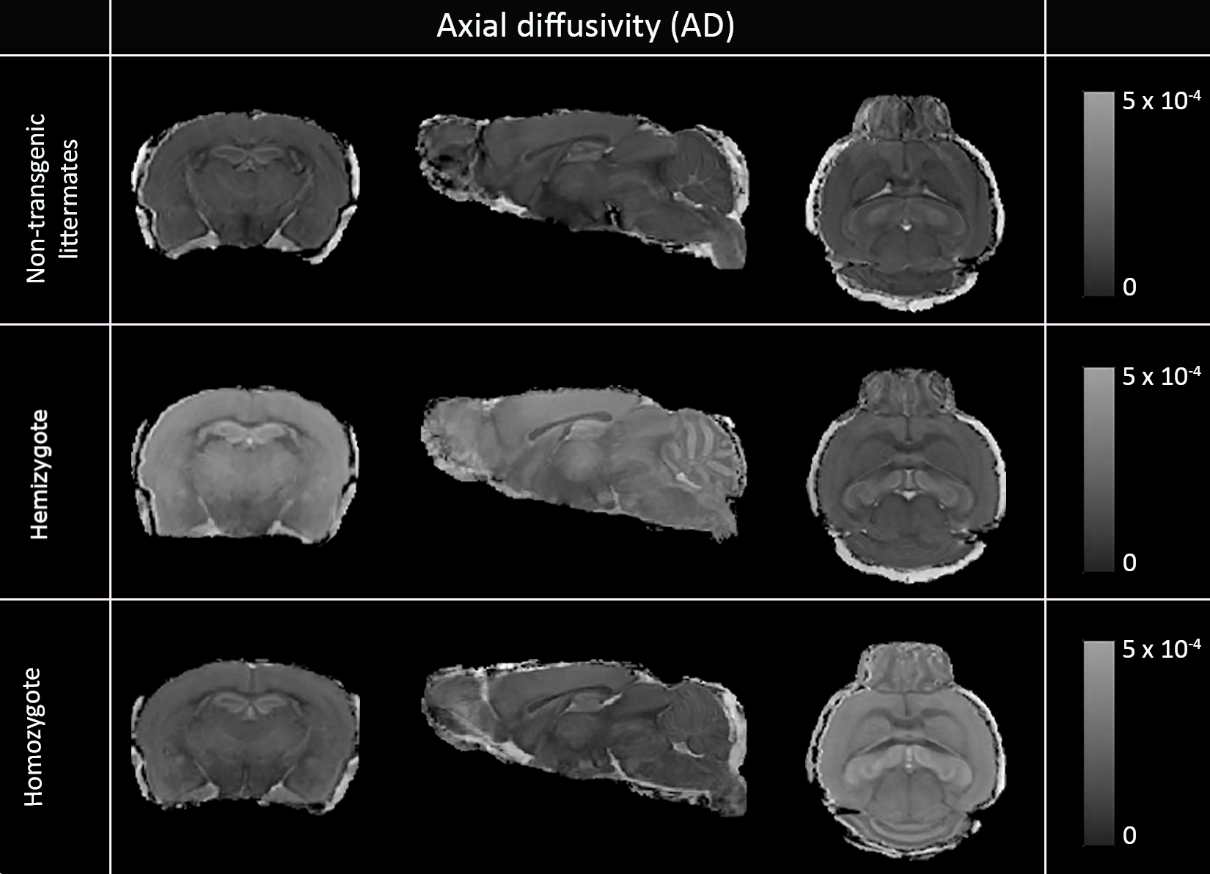

SFig 4

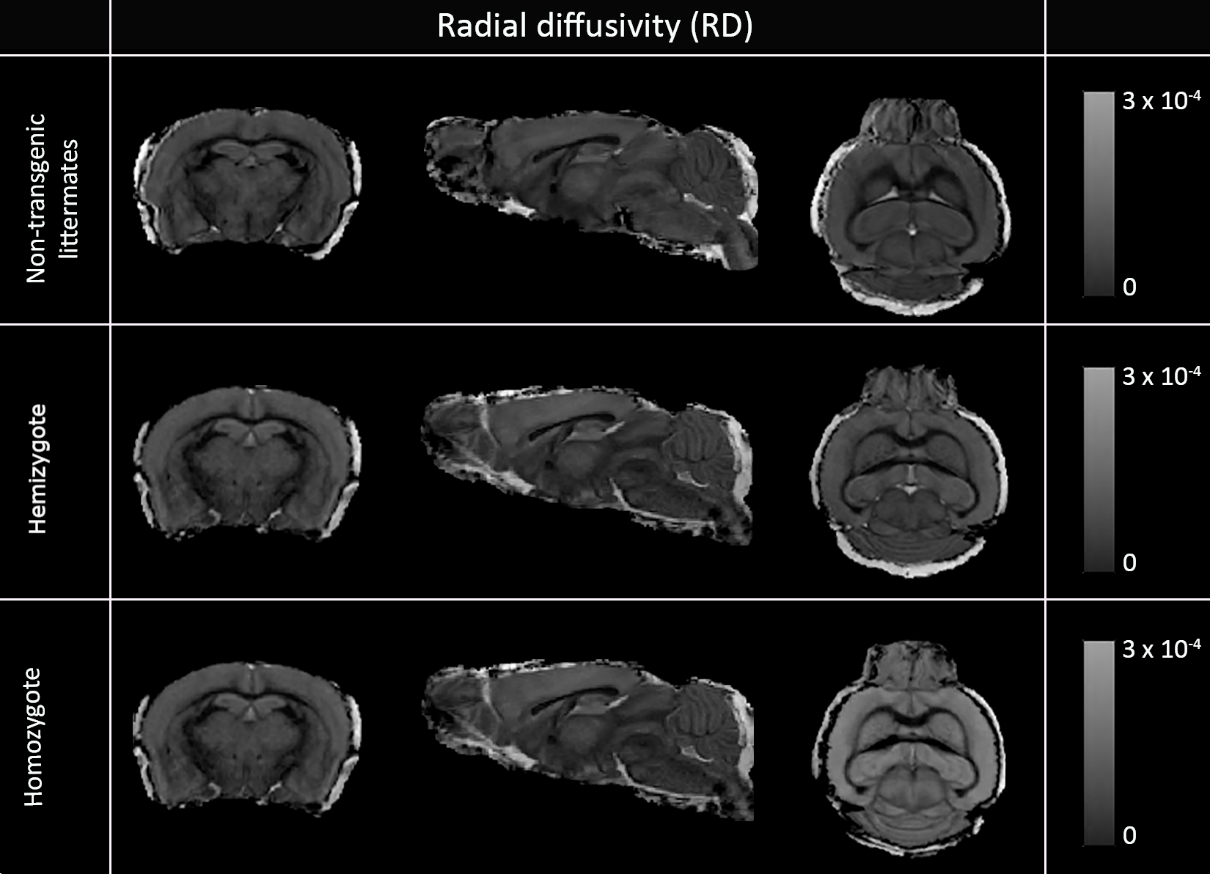
